# Doctoring Direct-to-Consumer Genetic Tests with DNA Spike-Ins

**DOI:** 10.1101/2022.04.01.486752

**Authors:** Peter Ney, Arkaprabha Bhattacharya, David Ward, Luis Ceze, Tadayoshi Kohno, Jeff Nivala

## Abstract

Direct-to-consumer (DTC) genetic testing companies have provided personal genotyping services to millions of customers. Customers mail saliva samples to DTC service providers to have their genotypes analyzed and receive back their raw genetic data. Both consumers and the DTC companies use the results to perform ancestry analyses, relative matching, trait prediction, and estimate predisposition to disease, often relying on genetic databases composed of the data from millions of other DTC-genotyped individuals. While the digital integrity risks to this type of data have been explored, we considered whether data integrity issues could manifest upstream of data generation through physical manipulation of DNA samples themselves, for example by adding synthetic DNA to a saliva sample (“spiked samples”) prior to sample processing by a DTC company. Here, we investigated the feasibility of this scenario within the standard DTC genetic testing pipeline. Starting with the purchase of off-the-shelf DTC genetic testing kits, we found that synthetic DNA can be used to precisely manipulate the results of saliva samples genotyped by a popular DTC genetic testing service and that this method can be used to modify arbitrary single nucleotide polymorphisms (SNPs) in multiplex to create customized doctored genetic profiles. This capability has implications for the use of DTC-generated results and the outcomes of their downstream analyses.

## Main Text

Since the advent of high-density genotyping and next-generation DNA sequencing, there has been tremendous growth in the amount of genetic data that is collected, processed, and stored (*1–3*). Some of the biggest producers of genetic data have been low-cost consumer facing genotyping services, so called direct-to-consumer (DTC), that process samples from the general public using high-density genotyping arrays. DTC companies use this data to give customers insights into their ancestry, find close relatives, predict traits, and find predisposition to disease (*4,5*). Presently, millions of customers have been genotyped via DTC testing providers, with the largest DTC providers having processed and stored data from over 10 million individuals (*6*).

Given this scale, there is a lot at stake in the design and implementation of the DTC genomics ecosystem. Any genotypes or metadata stored in DTC databases are at risk of unauthorized access or data theft (*7,8*). In addition to typical cybersecurity concerns, like data breaches, poorly designed analysis tools and visualizations specific to DTC genomics have also been shown to leak private genetic information (*9–11*). DTC design not only affects those that are tested but can impact close relations or broader society. Genetic genealogy databases have repeatedly been shown to be sufficient to infer the identity of unknown DNA samples or data — including criminal forensic samples and anonymous research subjects (*12–14*).

In this work, we consider a different aspect of genetic testing risk that has received less attention: the underlying assumption that genetic results stored in DTC databases are genuine and could not have not been doctored in an adversarial manner prior to the data generation process. This is especially important in the DTC industry because there is extensive digital data sharing by individuals to third-parties and, most relevant to this work, there is no sample provenance (*4*). Samples are collected and submitted to the DTC services directly by the general public with minimal sample verification. This means that essentially anyone with moderate technical means could submit anonymous, forged, or pseudoanonymous saliva samples directly to a DTC genetic testing service and obtain apparently authentic genetic results. This data could then ultimately be used in downstream analyses (Fig. 1).

**Fig 1.**
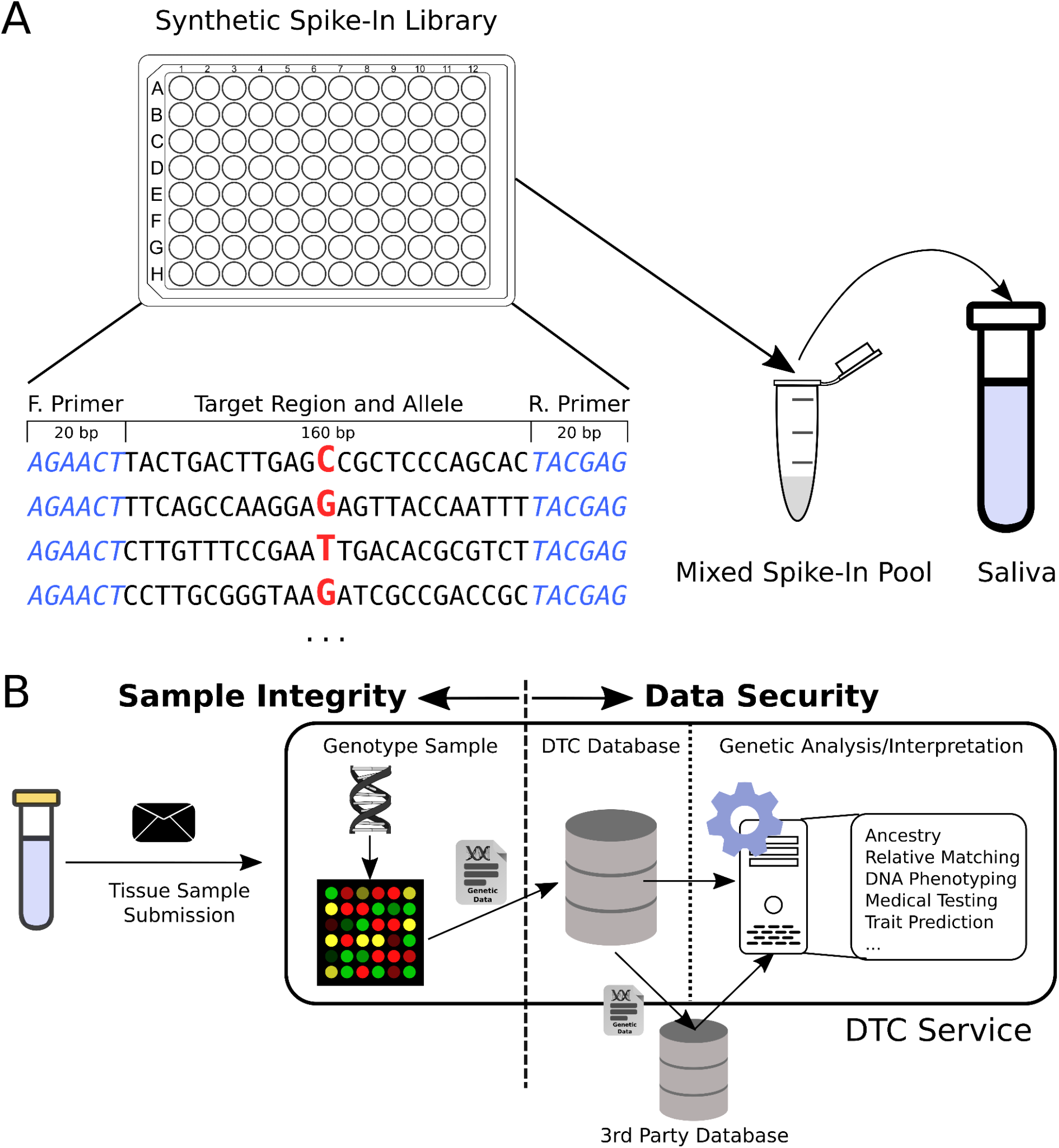
DNA spike-ins and possible effects on the DTC ecosystem. (**A**) Synthetic DNA strands from a preconfigured spike-in library are selected, pooled, and spiked into a saliva to forge a desired genotype. (**B**) The spiked-in tissue sample is sent to a DTC service for genotyping or sequencing. The left side of the dotted line denotes the phase where genetic data is in a physical form and contains no integrity checks. On the right portion of the figure genetic information is shared and processed as digital information, raising typical data security concerns.

The most popular DTC genomic testing technologies are DNA microarray machines (e.g., Illumina’s Global Screening Array), which are capable of characterizing hundreds of thousands of sites of genetic variation, so called single nucleotide polymorphisms (SNPs), that are known to vary throughout the human population according to ancestry and contribute to a range of phenotypes, from appearance to disease predisposition (*15,16*). Thus, the ability to manipulate microarray results would make it possible to generate, for example, customized physical phenotype profiles. Motivated by this possibility, we decided to explore the potential for manipulating microarray results generated by a DTC genetic testing service at the physical level.

Here, we show that synthetic DNA can be mixed with saliva to manipulate the results of otherwise natural consumer samples genotyped by a DTC service. We find that simply mixing negligible volumes of specifically designed synthetic DNA into a saliva sample is sufficient to arbitrarily modify SNPs to a desired genotype and that different synthetic DNA strands can be combined in an additive manner to change at least dozens of individual SNPs simultaneously. These altered genotypes led to different phenotypic predictions by the DTC service, demonstrating that physical sample spike-ins can alter downstream interpretation in commercial pipelines.

To develop this methodology, our first objective was to create a simple saliva spiking protocol that minimized prepwork and would allow sets of target SNPs to be modified arbitrarily by people with minimal resources and expertise. We began with a synthetic DNA construct design that could be used to target specific SNP locations in the human genome. In this design, each synthetic DNA strand was 200-bp in length with the middle 160-bp encoding for the human genome reference sequence (GRCh37) surrounding a particular SNP. Invariant 20-bp sequences were also included on each end of the DNA construct as universal PCR priming sites for fragment amplification. The middle nucleotide of each strand was then used to encode for the desired SNP. Importantly, this construct design would make it possible to forge multiple SNPs at once, as the desired fragments could be pooled together before mixing into the saliva sample.

To test this design, we constructed a 48-SNP spoofing library with two possible alleles per SNP, for a total of 96 strands. We chose SNPs with previously known correlations to physical traits or appearance (Supplement Table S1). The ssDNA library was ordered using a commercial oligo service and each strand amplified into dsDNA via PCR (Supplement Table S2). Since the middle 160-bp of each strand in the library is complementary to the adjacent genomic sequence nearby the SNP, any SNPs within +/− 80-bp of each other will have overlapping sequences and could potentially cross-react with microarray probe binding. To understand this effect, 8 of the 40 SNPs in the library were specifically chosen to be overlapping.

In the subsequent experiments described here, specific SNP fragments from this synthetic DNA library were spiked into saliva samples collected from two different individuals^1^ and submitted to a major DTC genetic testing service for genotyping. A total of 25 saliva samples from these two individuals were submitted between November 2020 - June 2021. These include 2 controls (no spike-in), 5 for protocol debugging/testing, 8 to test the effect of spike-in concentration, and 10 to explore multiplex SNP spike-ins. To minimize the impact on the DTC company, we chose options that excluded the samples from research studies and relative matching functionality.

We started with the simplest case of forging a single SNP (rs17822931) in saliva from one individual at various concentrations. The individual was naturally homozygous CC at this locus (determined from the control sample with no spike-in DNA). We used a spike-in strand encoding the T allele. In this experiment, there are four possible genotype outcomes: no change (CC), single base flip (CT), double base flip (TT), or no call (--). We consider a SNP successfully modified if the SNP is called and the genotype has at least one base flip towards the synthetic allele (i.e., CT or TT in this example). Note, if the individual was heterozygous CT then only a single base flip is possible — TT. The synthetic T-allele fragment for rs17822931 was serially diluted and spiked-in at a wide range of DNA quantities (1000 ng - 0.01 ng) in 8 unique saliva samples for the same individual and mailed to the DTC company (Supplementary Table S3A). After several weeks, we received the genotyping results for each of these samples from the company. As anticipated, at higher spike-in quantities (> 10 ng) the SNP was flipped to homozygous TT, intermediate quantities (0.1-1 ng) the SNP was either partially flipped (CT) or was a no call, and at low quantities (< 0.01 ng) there was no modification to the natural SNP (CC). These results show that synthetic DNA can be used to manipulate DTC genotyping outcomes. It also suggests that spike-in DNA concentration may be adjusted to provide either homozygous or heterozygous results.

Next, we performed multiplexed SNP forging experiments by pooling multiple synthetic strands for different SNPs into each saliva sample. This was tested across ten samples using saliva from both individuals. Each sample was spiked with between 20-48 strands from the synthetic SNP library at three dilutions (1X, 0.1X, 0.01X); The first individual was spiked with 20 SNPs and the second individual with 39 or 48 SNPs (Supplementary Table S3B). The spoofed allele of each SNP was chosen to be different from the genotype at that locus for each individual; when naturally heterozygous, a homozygous allele was arbitrarily chosen.

After analysis using the DTC service, we found that the synthetic DNA spike-ins altered the allele of the targeted SNP in all cases when a genotype was called. Specifically, we observed that the genotypes for the 10 samples were split between completely successful modifications (45.5%; 155/341), i.e., the genotype was modified to the desired homozygous result, no-calls (52.2%; 178/341), and a small number (2.3%; 8/341) were heterozygous calls with one base altered (Supplementary table S3C). The average no-call rate of 52.2% for the spiked-in SNPs is higher than the average overall no-call rate of 0.27%, indicating that the addition of synthetic DNA SNP fragments are lowering the quality of SNPs that are targeted. However, the spiked-in DNA did not seem to otherwise affect data quality for the non-targeted SNPs because there was little relationship between spike-in quantity and no call rate (r^2^ - 0.141) or miscall rate (r^2^ - 0.055), where a miscall is any SNP with a different genotype than the control sample (Supplementary Fig. S1).

For the 8 overlapping SNPs, we found that they had a particularly high no-call rate (89.6%; 43/48), likely due to interference between adjacent SNPs (i.e., within +/-80bp). Due to poor amplification, one of the SNPs (rs1800414) had an abnormally low concentration, and thus, was spiked-in at lower concentrations than the other SNPs. Similar to what was seen in the single SNP concentration experiment with lower concentrations, a heterozygous genotype was called. Of the remaining 39 SNPs, whenever a SNP was called (50.3% call rate) it was always homozygous with the spike-in allele, further indicating that the probes are likely saturated with the spike-in strands (outcompeting the natural genomic DNA fragments), resulting in a homozygous genotype when the call passes quality controls.

Taken together, these results indicate that a simple SNP spike-in, with non-adjacent SNPs and at sufficient concentration, can be pooled together to alter at least dozens of SNPs in a single sample with around 50% effectiveness. The partial effectiveness (high probability of no calls) would not be an issue in many contexts because most SNP-based interpretation tools manage no-calls gracefully, as they commonly occur in normal genotype results. To manage this, models simply drop the no-calls and rely on the subset of the SNPs in the model which have been called. To see how the DTC service would interpret the forged SNPs we relied on it’s built in phenotype prediction to see how the results were affected. While the underlying models were not public, we attempted to include SNPs in the spoofing library which are known to relate to tested phenotypes. These include markers for pigmentation (e.g., eye, hair, and skin color), facial features (e.g., ear lobe shape and cleft chin), and others (e.g., finger ratio, unibrow). Due to the underlying variability in flipping a given spike-in SNP, traits were forged only some of the time. However, for all but two of the traits (male hair loss and unibrow) all of the targeted traits were forged in at least some of the samples, with many samples having multiple forged traits (Table 1). We also found that the spike-in samples’ heritage prediction results were altered relative to the control sample. These results indicate that forging SNPs with publicly known phenotypic effects may be sufficient to change digital predictions offered by DTC services, highlighting the link between physical sample manipulation and possible digital impacts.

**Table 1.** Spike-In Effect on Predicted Traits. (**A**) Predicted traits reported by the DTC testing company for the 10 multiplex spike-in samples for the two tested individuals. The four samples for Individual 1 are denoted as samples M1-M4 and the six for Individual 2 as samples M5-M10. Key for observed phenotypes: cleft chin (yes/no), eye color (blue/light/dark), hair color (blond, brunette, red, dark), skin pigment (light-to-medium, dark), facial hair thickness (less thick, thicker), iris patterns (crypts, rings, and furrows), male hair loss (low chance), earlobe type (attached, unattached), earwax type (wet, dry), digit ratio (index finger longer, ring finger longer), freckles (yes, no), hair thickness (average, thicker than average), hair type (wavy, straight), unibrow (no). (**B**) Raw spike-in effect on the 22 SNPs altered in Individual 1 (samples M1-M4). Two dashes (--) indicate a no call and “No spike” indicates that specific SNP was not spiked into that sample.

|  | Individual 1 |  |  |  |  | Individual 2 |  |  |  |  |  |  |
| --- | --- | --- | --- | --- | --- | --- | --- | --- | --- | --- | --- | --- |
|  | Orig | M1 | M2 | M3 | M4 | Orig | M5 | M6 | M7 | M8 | M9 | M10 |
| **Cleft Chin** | Yes | No | No | No | No | No | No | Yes | Yes | No | Yes | Yes |
| **Eye Color** | Dark | Dark | Dark | Dark | Dark | Dark | Blue | Dark | Blue | Light | Blue | Dark |
| **Hair Color** | Dark | Dark | Dark | Dark | Dark | Brown | Red | Red | Blond | Red | Red | Brown |
| **Skin Pigment** | Light / Med | Dark | Dark | Light / Med | Light / Med | Light / Med | Light / Med | Light / Med | Light / Med | Light / Med | Light / Med | Light / Med |
| **Facial Hair Thickness** | Less | More | More | Less | More | Less | Less | Less | Less | Less | More | Less |
| **Iris Patterns** | C,R | C,R | C,R,F | C,R | C,R,F | C,R,F | C,R | C,R | C,F | C,R,F | C | C,R |
| **Male Hair Loss** | Low | Low | Low | Low | Low | Low | Low | Low | Low | Low | Low | Low |
| **Earlobe Type** | Unatt. | Attach | Attach | Attach | Attach | Attach | Unatt. | Attach | Unatt. | Attach | Unatt. | Attach |
| **Earwax Type** | Wet | Dry | Dry | Dry | Dry | Wet | Wet | Wet | Wet | Wet | Wet | Wet |
| **Digit Ratio** | Ring | Ring | Index | Ring | Ring | Index | Index | Index | Ring | Index | Ring | Ring |
| **Freckles** | No | No | No | No | No | No | Yes | No | Yes | Yes | Yes | No |
| **Hair Thickness** | Avg | Avg | Thick | Avg | Thick | Avg | Avg | Avg | Avg | Avg | Thick | Avg |
| **Hair Type** | Wavy | Wavy | Str. | Wavy | Str. | Wavy | Str. | Wavy | Wavy | Str. | Str. | Str. |
| **Unibrow** | No | No | No | No | No | No | No | No | No | No | No | No |

**Table 1.** Spike-In Effect on Predicted Traits.
| rsid | Trait | Original Genotype | Spike-in Allele | M1 | M2 | M3 | M4 |
| --- | --- | --- | --- | --- | --- | --- | --- |
| rs11684042 | Cleft Chin | AG | G | GG | GG | GG | GG |
| rs4864809 | Facial Hair Thickness | GG | A | AA | AA | -- | AA |
| rs4900109 | Iris Pattern | TG | G | GG | GG | GG | GG |
| rs10235789 |  | CC | T | -- | -- | -- | -- |
| rs3739070 |  | AC | A | -- | AA | -- | AA |
| rs2080401 | Earlobe Type | AA | C | CC | CC | CC | CC |
| rs17822931 | Earwax Type | CC | T | TT | TT | TT | TT |
| rs2395845 | Unibrow | CC | A | AA | AA | AA | AA |
| rs756853 | Male Hair Loss | AA | G | -- | GG | -- | GG |
| rs2180439 |  | TC | T | TT | TT | -- | TT |
| rs2497938 |  | TT | C | CC | CC | CC | CC |
| rs11158820 | Digit Ratio | AG | A | AA | AA | AA | AA |
| rs2332175 |  | AG | A | AA | AA | -- | AA |
| rs314277 |  | CC | A | -- | AA | -- | -- |
| rs3827760 | Hair Thickness/Type | AA | G | -- | GG | -- | GG |
| rs11803731 | Hair Type | AA | T | TT | TT | -- | TT |
| rs17646946 |  | GG | A | AA | AA | AA | AA |
| rs7349332 |  | CC | T | TT | TT | TT | -- |
| rs12896399 | Eye Color, Hair Color, Skin Color, Freckles | TG | T | No Spike | No Spike | -- | TT |
| rs1470608 |  | TG | T | -- | TT | No Spike | No Spike |
| rs12913832 |  | AG | G |  | N/A | -- | -- |
| rs1426654 |  | AA | G | GG | GG | No Spike | No Spike |

In summary, we have described here the first demonstration, to our knowledge, of how the DTC genetic testing pipeline can be doctored at the physical level. Data integrity issues can affect all stages of the DTC pipeline from sample processing to storage, analysis, and data sharing. Cryptographic techniques like digital signature schemes are a good approach to ensure the authenticity of digital data, but these results highlight the need to consider the integrity of the physical sample channel as well (*12*). Whenever DNA is processed in potentially adversarial environments it may be worthwhile to confirm that samples do not contain synthetic DNA (e.g., methylation analysis), as has been suggested in other genetic contexts, like forensics (*17*). While the emphasis of this work was on the DTC testing ecosystem, we believe there are broader lessons for the larger sequencing industry. Most data processing equipment and algorithms are likely not robust to adversarial manipulation because they are only designed to tolerate naturally occurring noise. Therefore, the vulnerabilities raised in this work are likely to impact most types of genotyping, sequencing, and associated analysis pipelines.

## Methods

### Synthetic SNP fragment preparation

Single-stranded DNA oligonucleotides were ordered from Integrated DNA Technologies (IDT). To prepare oligos for spiking experiments, each oligo was amplified by PCR (KAPA) using the universal primers, purified with AMPure XP magnetic beads (Beckman), and quantified with qPCR prior to spiking experiments.

### DTC saliva genotyping kit spike-in

DTC saliva genotyping kits were ordered from a commercial vendor. Saliva samples were submitted to the DTC company according to the recommended protocol except with the additional step of adding the desired synthetic SNP DNA fragment(s) at the appropriate concentrations. Water was used to dilute the synthetic DNA stocks prior to addition to the saliva sample when required.

## Acknowledgments

We thank members of the UW Molecular Information Systems Lab and the UW Security and Privacy Research Lab for feedback on this work.

## Author contributions

Conceptualization: PN, LC, TK, JN
Methodology: PN, AB, DW, LC, TK, JN
Investigation: PN, AB, DW, JN
Visualization: PN
Funding acquisition: PN, LC, TK, JN
Supervision: TK, JN
Writing – original draft: PN, JN
Writing – review & editing: PN, TK, JN

## Data and materials availability

Raw data for the targeted SNPs are included in the supplement. Release of full genotype files will be considered upon request.

## Footnotes

1 The two individuals were co-authors of this work. The DTC Terms-of-Service required that submitted saliva come from the person registering the sample or with their consent.

