## Supplementary Materials for "Doctoring Direct-to-Consumer Genetic Tests with DNA Spike-Ins"

[1]Paul G. Allen School of Computer Science & Engineering. University of Washington; Seattle, Washington, USA.

[2]Molecular Engineering and Sciences Institute. University of Washington; Seattle, Washington, USA.

**A**

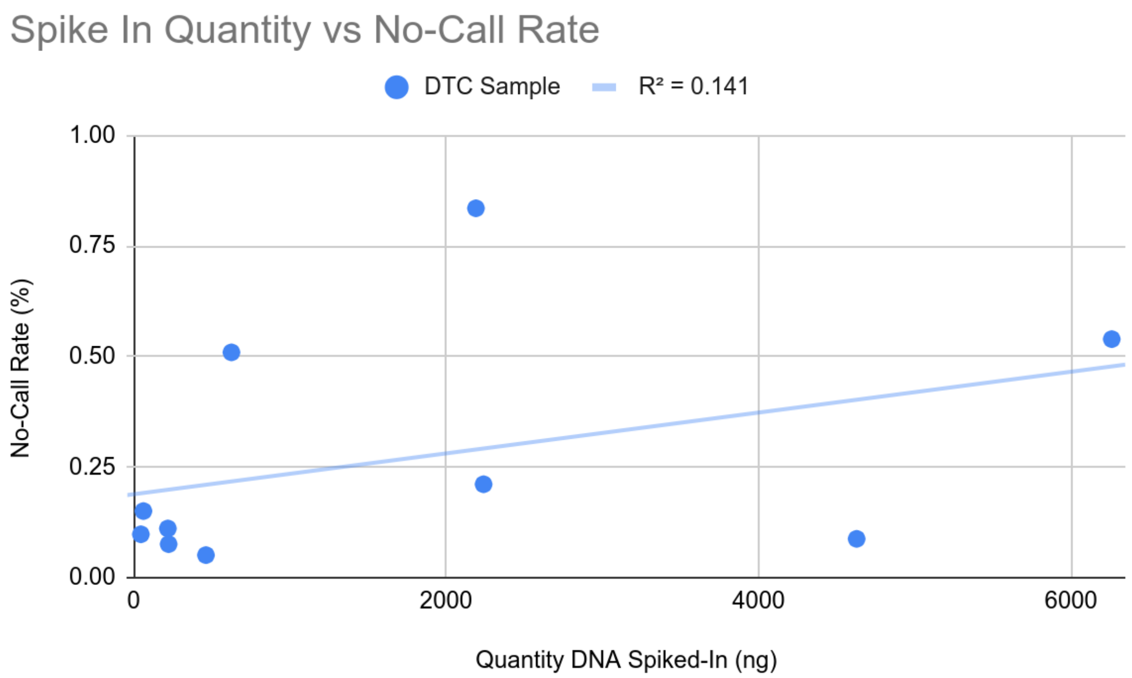


**B**

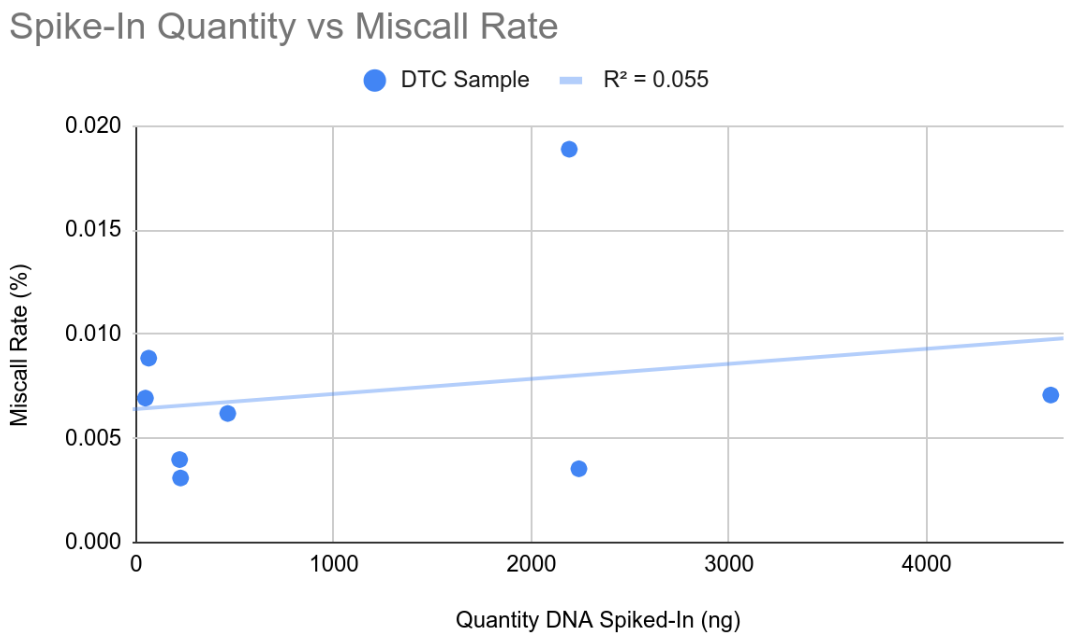


**Figure S1.** Spike-in effect on overall run quality. Weak association between the total quantity of spike-in DNA (ng) and (**A**) overall no-call rate (%) and (**B**) overall miscall rate (%). SNP calls are considered miscalls if the call differs from the control sample.

| Trait Type | SNPs (rssid) |
|---|---|
| Cleft Chin | rs11684042 |
| Digit Ratio | rs11158820, rs2332175, rs314277 |
| Ear Wax Type | rs17822931 |
| Earlobe Type | rs2080401 |
| Eye Color, Hair Color, Skin Pigmentation, Freckles | rs16891982, rs12203592, rs4959270, rs683, rs10756819, rs1042602, rs1393350, rs1126809, rs12821256, rs12896399, rs1545397, rs1800414, rs1800407, rs1470608, rs1129038, rs12913832, rs1667394, rs1426654, rs3114908, rs1805005, rs1805006, rs2228479, rs11547464, rs1805007, rs201326893, rs1110400, rs1805008, rs885479, rs1805009, rs6059655 |
| Facial Hair Thickness | rs4864809 |
| Hair Thickness | rs3827760 |
| Hair Type | rs3827760, rs11803731, rs17646946, rs7349332 |
| Iris Patterns | rs4900109, rs10235789, rs3739070 |
| Male Hair Loss | rs756853, rs2180439, rs2497938 |
| Unibrow | rs2395845 |

**Table S1.** Targeted traits and corresponding SNPs. The eye color, hair color, skin pigmentation, and freckles SNPs were selected from the HirisPlex System (*18).*

| rsid | Chr:Pos | Strand-1 | | Strand-2 | |
|---|---|---|---|---|---|
| | | Allele | DNA conc. (ng/µL) | Allele | DNA conc. (ng/µL) |
| rs17646946 | 1:152062767 | A | 96.3 | G | 106.1 |
| rs11803731 | 1:152083325 | A | 92.8 | T | 65 |
| rs3827760 | 2:109513601 | A | 71.8 | G | 71.1 |
| rs11684042 | 2:145684278 | A | 120.9 | G | 101.3 |
| rs2080401 | 2:171540823 | A | 61.1 | C | 76 |
| rs7349332 | 2:219756383 | C | 158 | T | 148 |
| rs2395845 | 2:222993315 | A | 125.7 | C | 199 |
| rs3739070 | 2:239306268 | A | 124.1 | C | 174 |
| rs4864809 | 4:54511913 | A | 151 | G | 179 |
| rs16891982 | 5:33951693 | C | 151 | G | 137 |
| rs12203592 | 6:396321 | C | 155 | T | 147 |
| rs4959270 | 6:457748 | A | 135 | C | 123.7 |
| rs314277 | 6:105407662 | A | 101.2 | C | 78.2 |
| rs756853 | 7:18890000 | A | 67.1 | G | 80.9 |
| rs10235789 | 7:83656800 | C | 88.8 | T | 103.8 |
| rs683 | 9:12709305 | A | 104.3 | C | 106.4 |
| rs10756819 | 9:16858084 | A | 81.4 | G | 75.5 |
| rs1042602 | 11:88911696 | A | 71.8 | C | 72.3 |
| rs1393350 | 11:89011046 | A | 90.4 | G | 87.8 |
| rs1126809 | 11:89017961 | A | 56.1 | G | 72.7 |
| rs12821256 | 12:89328335 | C | 36.4 | T | 61.8 |
| rs2332175 | 14:70345411 | A | 57.4 | G | 40.9 |
| rs11158820 | 14:70347348 | A | 30.1 | G | 43.2 |
| rs4900109 | 14:92763391 | G | 49 | T | 69.8 |
| rs12896399 | 14:92773663 | G | 75.1 | T | 160.7 |
| rs1545397 | 15:28187772 | A | 134.5 | T | 140.4 |
| rs1800414 | 15:28197037 | C | 0.8 | T | 167.2 |
| rs1800407 | 15:28230318 | C | 144.2 | T | 120.3 |
| rs1470608 | 15:28288121 | G | 152.2 | T | 172.9 |
| rs1129038 | 15:28356859 | C | 119.2 | T | 95 |
| rs12913832 | 15:28365618 | A | 213.5 | G | 139.4 |
| rs1667394 | 15:28530182 | C | 191.1 | T | 218.6 |

| | | | | | |
|---|---|---|---|---|---|
| rs1426654 | 15:48426484 | A | 200.7 | G | 176 |
| rs17822931 | 16:48258198 | C | 137.3 | T | 128.5 |
| rs3114908 | 16:89383725 | C | 177.1 | T | 155.8 |
| rs1805005 | 16:89985844 | G | 94.6 | T | 106.8 |
| rs1805006 | 16:89985918 | A | 258.1 | C | 193.1 |
| rs2228479 | 16:89985940 | A | 200.4 | G | 265.9 |
| rs11547464 | 16:89986091 | A | 192.1 | G | 166.2 |
| rs1805007 | 16:89986117 | C | 149.4 | T | 202.8 |
| rs201326893 | 16:89986122 | A | 199.2 | C | 195 |
| rs1110400 | 16:89986130 | C | 125.6 | T | 151.2 |
| rs1805008 | 16:89986144 | C | 152.3 | T | 179.4 |
| rs885479 | 16:89986154 | A | 204.9 | G | 202.6 |
| rs1805009 | 16:89986546 | C | 180.8 | G | 181.1 |
| rs2180439 | 20:21853100 | C | 179.1 | T | 221.4 |
| rs6059655 | 20:32665748 | A | 195.5 | G | 255.8 |
| rs2497938 | X:66563018 | C | 160 | T | 123.7 |

**Table S2** Spike-In library. 48-SNP library with 2 alleles per SNP. Genomic positions use GRCh37.

**A**

| Sample ID | Spike-in Mass (ng DNA) | Genotype (rs17822931) |
|---|---|---|
| WT | 0 | CC |
| C1 | 0.001 | CC |
| C2 | 0.01 | CC |
| C3 | 0.1 | CT |
| C4 | 1 | No Call |
| C5 | 10 | TT |
| C6 | 100 | TT |
| C7 | 500 | TT |
| C8 | 1000 | TT |

**B**

| Sample ID | Individual | Spike-In SNPs | Number Flipped | Ambiguous | | Flip Rate | Overlapping | Dilution Factor |
|---|---|---|---|---|---|---|---|---|
| | | | | No-call | Heterozygous | | | |
| M1 | 1 | 20 | 16 | 6 | 0 | 72.7 | 0 | 1x |
| M2 | 1 | 20 | 19 | 1 | 0 | 95.0 | 0 | 0.1x |
| M3 | 1 | 20 | 9 | 11 | 0 | 45.0 | 0 | 1x |
| M4 | 1 | 20 | 16 | 4 | 0 | 80.0 | 0 | 0.1x |
| M5 | 2 | 39 | 11 | 28 | 0 | 28.2 | 0 | 1x |
| M6 | 2 | 39 | 30 | 8 | 1 | 76.9 | 0 | 0.1x |
| M7 | 2 | 39 | 20 | 18 | 1 | 51.3 | 0 | 0.01x |
| M8 | 2 | 48 | 14 | 25 | 0 | 35.9 | 9 | 1x |
| M9 | 2 | 48 | 11 | 27 | 1 | 28.2 | 9 | 0.1x |
| M10 | 2 | 48 | 11 | 28 | 0 | 28.2 | 9 | 0.01x |

## C

| Individual 1 | | | | | | | | |
|---|---|---|---|---|---|---|---|---|
| Number | **SNP ID** | **Chr:Pos** | **Genotype** | **Spike-in Allele** | **M1** | **M2** | **M3** | **M4** |
| 1 | rs17646946 | 1:152062767 | GG | A | AA | AA | AA | AA |
| 2 | rs11803731 | 1:152083325 | AA | T | TT | TT | -- | TT |
| 3 | rs3827760 | 2:109513601 | AA | G | -- | GG | -- | GG |
| 4 | rs11684042 | 2:145684278 | AG | G | GG | GG | GG | GG |
| 5 | rs2080401 | 2:171540823 | AA | C | CC | CC | CC | CC |
| 6 | rs7349332 | 2:219756383 | CC | T | TT | TT | TT | -- |
| 7 | rs2395845 | 2:222993315 | CC | A | AA | AA | AA | AA |
| 8 | rs3739070 | 2:239306268 | AC | A | -- | AA | -- | AA |
| 9 | rs4864809 | 4:54511913 | GG | A | AA | AA | -- | AA |
| 10 | rs314277 | 6:105407662 | CC | A | -- | AA | -- | -- |
| 11 | rs756853 | 7:18890000 | AA | G | -- | GG | -- | GG |
| 12 | rs10235789 | 7:83656800 | CC | T | -- | -- | -- | -- |
| 13 | rs2332175 | 14:70345411 | AG | A | AA | AA | -- | AA |
| 14 | rs11158820 | 14:70347348 | AG | A | AA | AA | AA | AA |
| 15 | rs4900109 | 14:92763391 | TG | G | GG | GG | GG | GG |
| 16 | rs12896399 | 14:92773663 | TG | T | N/A | N/A | -- | TT |
| 17 | rs1470608 | 15:28288121 | TG | T | -- | TT | N/A | N/A |
| 18 | rs12913832 | 15:28365618 | AG | G | N/A | N/A | -- | -- |
| 19 | rs1426654 | 15:48426484 | AA | G | GG | GG | N/A | N/A |
| 20 | rs17822931 | 16:48258198 | CC | T | TT | TT | TT | TT |
| 21 | rs2180439 | 20:21853100 | TC | T | TT | TT | -- | TT |
| 22 | rs2497938 | X:66563018 | TT | C | CC | CC | CC | CC |

| Individual 2 | | | | | | | | | | |
|---|---|---|---|---|---|---|---|---|---|---|
| Number | SNP ID | Chr:Pos | Original Genotype | Spike-in Allele | M5 | M6 | M7 | M8 | M9 | M10 |
| 1 | rs17646946 | 1:152062767 | GG | A | AA | -- | -- | AA | AA | AA |
| 2 | rs11803731 | 1:152083325 | AA | T | -- | -- | -- | -- | TT | TT |
| 3 | rs3827760 | 2:109513601 | AA | G | -- | -- | -- | -- | GG | -- |
| 4 | rs11684042 | 2:145684278 | GG | A | -- | AA | AA | -- | AA | AA |
| 5 | rs2080401 | 2:171540823 | AC | A | AA | -- | AA | -- | AA | -- |
| 6 | rs7349332 | 2:219756383 | TC | C | -- | CC | -- | -- | CC | -- |
| 7 | rs2395845 | 2:222993315 | CC | A | AA | AA | -- | AA | AA | -- |
| 8 | rs3739070 | 2:239306268 | AA | C | CC | CC | -- | -- | CC | CC |
| 9 | rs4864809 | 4:54511913 | GG | A | -- | -- | -- | -- | AA | -- |
| 10 | rs16891982 | 5:33951693 | GG | C | -- | -- | -- | -- | -- | -- |
| 11 | rs12203592 | 6:396321 | CC | T | TT | -- | -- | TT | TT | -- |
| 12 | rs4959270 | 6:457748 | CC | A | -- | -- | -- | -- | AA | -- |
| 13 | rs314277 | 6:105407662 | AC | C | -- | -- | CC | -- | CC | CC |
| 14 | rs756853 | 7:18890000 | AG | G | -- | -- | GG | -- | GG | GG |
| 15 | rs10235789 | 7:83656800 | TC | T | -- | -- | -- | -- | -- | -- |
| 16 | rs683 | 9:12709305 | AC | A | -- | -- | AA | -- | AA | AA |
| 17 | rs10756819 | 9:16858084 | AA | G | -- | GG | -- | -- | GG | GG |
| 18 | rs1042602 | 11:88911696 | CC | A | -- | -- | -- | -- | -- | -- |
| 19 | rs1393350 | 11:89011046 | GG | A | AA | -- | -- | -- | AA | AA |
| 20 | rs1126809 | 11:89017961 | GG | A | -- | -- | AA | -- | AA | AA |
| 21 | rs12821256 | 12:89328335 | TT | C | -- | -- | -- | -- | -- | -- |
| 22 | rs2332175 | 14:70345411 | AG | G | -- | -- | -- | -- | -- | -- |
| 23 | rs11158820 | 14:70347348 | AG | G | GG | GG | GG | GG | GG | GG |
| 24 | rs4900109 | 14:92763391 | TG | T | -- | -- | TT | -- | TT | -- |
| 25 | rs12896399 | 14:92773663 | TG | T | -- | -- | -- | -- | TT | TT |
| 26 | rs1545397 | 15:28187772 | AA | T | -- | -- | -- | -- | -- | -- |

| | | | | | | | | | | |
|---|---|---|---|---|---|---|---|---|---|---|
| 27 | rs1800414 | 15:28197037 | TT | C | -- | TC | -- | -- | TC | TC |
| 28 | rs1800407 | 15:28230318 | TC | C | CC | CC | -- | CC | CC | CC |
| 29 | rs1470608 | 15:28288121 | GG | T | -- | -- | -- | -- | TT | TT |
| 30 | rs1129038 | 15:28356859 | TC | T | -- | -- | TT | -- | TT | TT |
| 31 | rs12913832 | 15:28365618 | AG | G | GG | -- | -- | GG | GG | -- |
| 32 | rs1667394 | 15:28530182 | TC | T | -- | -- | TT | -- | TT | TT |
| 33 | rs1426654 | 15:48426484 | AA | G | GG | GG | -- | GG | GG | GG |
| 34 | rs17822931 | 16:48258198 | TC | C | -- | CC | -- | -- | -- | -- |
| 35 | rs3114908 | 16:89383725 | TC | C | CC | -- | -- | CC | -- | -- |
| 36 | rs1805005 | 16:89985844 | GG | T | -- | -- | -- | N/A | N/A | N/A |
| 37 | rs1805006 | 16:89985918 | CC | A | -- | -- | -- | N/A | N/A | N/A |
| 38 | rs2228479 | 16:89985940 | GG | A | -- | -- | -- | N/A | N/A | N/A |
| 39 | rs11547464 | 16:89986091 | GG | A | -- | -- | -- | N/A | N/A | N/A |
| 40 | rs1805007 | 16:89986117 | TC | C | -- | -- | -- | N/A | N/A | N/A |
| 41 | rs201326893 | 16:89986122 | CC | A | -- | -- | -- | N/A | N/A | N/A |
| 42 | rs1110400 | 16:89986130 | TT | C | -- | -- | -- | N/A | N/A | N/A |
| 43 | rs1805008 | 16:89986144 | CC | T | TC | -- | TC | N/A | N/A | N/A |
| 44 | rs885479 | 16:89986154 | GG | A | AG | AG | AG | N/A | N/A | N/A |
| 45 | rs1805009 | 16:89986546 | GG | C | CC | CC | -- | CC | CC | -- |
| 46 | rs2180439 | 20:21853100 | TC | T | TT | -- | -- | TT | TT | TT |
| 47 | rs6059655 | 20:32665748 | GG | A | -- | -- | -- | -- | AA | AA |
| 48 | rs2497938 | X:66563018 | TT | C | CC | CC | CC | CC | CC | CC |

**Table S3.** Spike-in experiment aggregate statistics and genotype results for targeted SNPs. (**A**) Serial dilution of spike-in strand targeting the A allele of rs17822931 into one individual. (**B**) Multiplex spike-in experiment summary statistics. (**C**) Genotypes for targeted SNPs in the multiplex spike-in experiments. Red genotypes indicate a heterozygous call. In Individual 2, samples M8-M10 were not spiked in with SNPs 36-44.

| Sample ID | Num Spike-In | Dilution Factor | Total Num SNPs | Num No-Calls | No-Call Rate (%) |
|---|---|---|---|---|---|
| M1 | 20 | 1X | 677857 | 1432 | 0.211 |
| M2 | 20 | 0.1X | 677857 | 514 | 0.076 |
| M3 | 20 | 1X | 677857 | 5663 | 0.835 |
| M4 | 20 | 0.1X | 677857 | 754 | 0.111 |
| M5 | 48 | 1X | 677857 | 3657 | 0.539 |
| M6 | 48 | 0.1X | 677857 | 3454 | 0.510 |
| M7 | 48 | 0.01X | 677857 | 1021 | 0.151 |
| M8 | 39 | 1X | 677857 | 595 | 0.088 |
| M9 | 39 | 0.1X | 677857 | 346 | 0.051 |
| M10 | 39 | 0.01X | 677857 | 665 | 0.098 |

**Table S4.** Quality statistics for the 10 multiple spike-in samples.
